# Effects of forest disturbance and regeneration on tree species composition and traits in DR Congo

**DOI:** 10.1101/2021.10.11.463162

**Authors:** Jonas Depecker, Anio Justin Asimonyio, Ronald Miteho, Yves Hatangi, Jean-Léon Kambale, Lauren Verleysen, Piet Stoffelen, Steven B. Janssens, Benoit Dhed’a, Filip Vandelook, Olivier Honnay

## Abstract

Despite their key role in biodiversity conservation, forests of the Congo Basin are increasingly threatened by human activities. Whereas deforestation is very conspicuous, it remains challenging to monitor the impact of forest degradation under a more or less intact canopy. Likewise, the outcomes of forest regeneration following agricultural abandonment remain poorly studied in the Congo Basin. Here, we surveyed 125 vegetation quadrats across 25 forest inventory plots in the Yangambi region, and based on historical land-use maps. We aimed to assess both the impact of anthropogenic disturbance and forest recovery on tree species diversity, tree functional traits, community composition, canopy cover, and forest structure, as compared to undisturbed old-growth forest. We found that undisturbed old-growth forest harboured more tree individuals, particularly individuals with a smaller diameter, as well as species with a higher wood density when compared to both disturbed old-growth and regrowth forest. Moreover, species diversity was highest in disturbed old-growth forest. Tree species composition was significantly different between undisturbed old-growth forest and the other two forest categories. Our findings indicate that anthropogenic disturbance results in a degraded forest structure and may reduce the forest’s ability to store carbon. Whereas species diversity recovered 60 years after agricultural abandonment, species composition remained different from undisturbed old-growth forest. Our study illustrates the need for more extensive vegetation surveys in the Congo Basin in order to quantify the effects of human disturbance under a closed canopy.

## 1. INTRODUCTION

Tropical and subtropical forests are by far the most floral rich biomes on earth (Kier et al., 2005). The African continent is home to the world’s second largest area of tropical forests, and 89% of them are located in the Congo Basin (Malhi et al., 2013). Whereas the African tropical forests appear to be less biodiverse than their Asian and American counterparts (Kier et al., 2005; Slik et al., 2015), Sosef et al. (2017) recorded 22,577 plant species in tropical African forests, of which 30% are endemic, but furthermore concluded that much botanical exploration remains to be done. In addition to their global importance for conserving biodiversity (Jung et al., 2021), tropical forests provide numerous ecosystem services including climate change mitigation by acting as natural carbon sinks (MEA, 2005). Tropical and subtropical forests globally account for 55% of gross carbon removals, with the relatively undisturbed forests of the Congo Basin removing up to six times more carbon than the forests in the Amazon region (Harris et al., 2021). Furthermore, the African carbon sink has been stable for the past three decades until 2015, while the Amazonian carbon sink has been declining since the 1990s (Hubau et al., 2020).

Despite their key role in biodiversity conservation and carbon sequestration, forests in tropical Africa are increasingly threatened by human activities (Tyukavina et al., 2018). Between 1990 and 2000, the annual net loss of forests corresponded to 3.3 million ha yr^−1^, increasing to 3.4 million ha yr^−1^ between 2000 and 2010, and further to 3.9 million ha yr^−1^ between 2010 and 2020 (FAO & UNEP, 2020). Africa is currently the only region where annual forest change rates are still increasing (FAO & UNEP, 2020). Deforestation on the African continent is mainly caused by small-scale subsistence agriculture (Jayathilake et al., 2021; Tegegne et al., 2016; Tyukavina et al., 2018), driven by very low agricultural yields (Carrasco et al., 2017; West et al., 2010) and a rapidly growing human population (UN, 2019).

Although deforestation is a very conspicuous and severe threat to biodiversity, degradation of tropical forests may even be more detrimental, based on the large spatial scales at which it occurs (Barlow et al., 2016). Forest degradation refers to within-forest disturbance and includes selective logging, removal of understorey vegetation, overexploitation of specific species, introduction of invasive species, and defaunation by hunting (Barlow et al., 2016; Sasaki et al., 2009; Tyukavina et al., 2018). Such degraded forests are commonly characterised by a significantly changed forest structure and species composition under a relatively intact canopy (Longo et al., 2016). In tropical Africa, forest degradation is primarily driven by selective logging, mainly for the small-scale production of charcoal which is still the most commonly used energy source in many Central African countries (Hosonuma et al., 2012; Jayathilake et al., 2021; Tyukavina et al., 2018). The degradation rate of forests in the Congo Basin has been estimated to be 31,7000 ha yr^−1^ between 2000 and 2005 (Ernst et al., 2013), whereas Lambin et al. (2003) estimated that the annual rate of forest degradation in Africa corresponds to approximately 50% of its annual rate of deforestation. This would imply that 1.95 million ha of African forest has been degraded between 2010 and 2020. However, Shapiro et al. (2021) recently reported that this assumption drastically underestimates the actual rate. Using a forest condition index, they estimated that between 2000 and 2016, over 23 million ha of forest has been degraded in the Congo Basin alone.

Although remote-sensing techniques are constantly improving, it remains challenging to monitor the extent and impact of forest degradation due to the interference with the often untouched canopy in degraded forests (Matricardi et al., 2020; Peres et al., 2006). Evaluation of the extent and impact of forest degradation must therefore largely rely on vegetation surveys on the ground (Heym et al., 2021). Whereas impacts of forest degradation on biodiversity have been frequently studied in the Neotropics (e.g. Carreño-Rocabado et al., 2012; Mirabel et al., 2020), Asia (e.g. Baboo et al., 2017; Millet et al., 2010), and other parts of Africa (e.g. Bobo et al., 2006; Oyugi et al., 2007), similar studies are rare in the Congo Basin (but see Hall et al., 2003; Ifo et al., 2016) and almost lacking in the Democratic Republic of Congo (DR Congo) (but see Makana & Thomas, 2006). Yet, as DR Congo harbours almost 60% of the African tropical forests (Malhi et al., 2013), it is essential to comprehend how plant community composition is affected by forest disturbance.

While conservation of existing tropical forests should still be the priority, the restoration of deforested land and degraded forests can aid in the recovery of tropical biodiversity and the recovery of ecosystem services (Brudvig, 2011; FAO & UNEP, 2020). Between 2000 and 2019, 0.7 million ha of forest has been actively or passively restored in Sub-Saharan Africa (FAO & UNEP, 2020). By 2030, the African Forest Landscape Restoration Initiative (AFR100) aims to increase the land surface under restoration to 100 million ha. Of this, DR Congo has committed to restore 8 million ha (AFR100, 2021). The biodiversity outcomes of restoration are governed by a combination of site-level factors, landscape-level factors, and historical contingencies (Brudvig, 2011), and; consequently, are difficult to predict (Novianti et al., 2018). However, during spontaneous forest regeneration, plant species richness and abundance tend to restore faster than species composition (Crouzeilles et al., 2016; Curran et al., 2014). Because of the high stochasticity and variability of restoration outcomes, more research is crucial to clear the path of restoration in different regions (Chazdon, 2008). Again, studies on biodiversity consequences of tropical forest regeneration are commonly found in the Neotropics (e.g. Letcher et al., 2009; Oberleitner et al., 2021) and tropical Asia (e.g. Hayward et al., 2021; Mahayani et al., 2020), but are rare in the Congo Basin (but see Bauters et al., 2019; Makelele et al., 2021). More specific, very little is known so far regarding the time that is needed for species composition to recover in this region.

Here, and based on historical land-use maps, we established a set of forest inventory plots in the Yangambi region in DR Congo aimed at quantifying both the impact of forest degradation and forest recovery following agricultural abandonment, on tree species composition and diversity. Based on 125 sampling quadrats across 25 forest inventory plots, we more specifically quantified differences among (i) disturbed old-growth forest, (ii) regrowth forest, and (iii) undisturbed old-growth forest in terms of tree species diversity, tree functional traits, community composition, forest structure, and canopy cover.

## 2. METHODS

### 2.1. Study site and set up

Vegetation sampling was conducted in the Yangambi region, in the Tshopo province in North-Eastern DR Congo, at approximately 100 km west of Kisangani. In Yangambi, the climate is characterised by two drier seasons (December-March & June-July) and two rainy seasons (April-May & August-November) (van Vliet et al., 2018). The general landscape type in this area is typical for the Congo Basin and is characterised by a mosaic of land tenures: the Yangambi Man and Biosphere Reserve created in 1979; the Ngazi Forest Reserve; a logging concession; and customary land (van Vliet et al., 2018). Prior to the Congolese independence in 1960, Yangambi housed a research station of INEAC (Institut National pour l’Etude Agronomique du Congo Belge), with multiple pastures for research and breeding activities. However, after INEAC changed into INERA (Institut National des Etudes et Recherches Agronomiques) in 1962, the activities at Yangambi started to diminish (Vanden Abeele et al., in press). As a result, many agricultural fields were abandoned, allowing forest to recover on these lands.

We established 25 forest inventory plots of 125 m × 125 m (1.56 ha) in an area of ca. 50 by 20 km, just North of the Congo River (Figure 1). Plots were assigned to three different forest categories by georeferencing (WGS84) them onto historical land-use maps which were issued by Evrard (1954). Plots located on historical agricultural land were classified as regrowth forest (R). Plots located in historically old-growth forests, adjacent to severely disturbed areas were considered as disturbed old-growth forest (DO). In all DO plots, there were clear signs of anthropogenic disturbance (e.g. selective logging and hunting, pers. obs.). The remaining plots that were located in historical old-growth forests were categorised as undisturbed old-growth forest (UO). Historically, access to these undisturbed sites was restricted by INEAC (Koy et al., 2019), while presently, there was no evidence of within-forest disturbance (pers. obs.) and local people confirmed that activities in these areas are still limited. This is furthermore confirmed by the occasional observation of large mammal species (e.g. *Smutsia gigantea* and *Tragelaphus eurycerus*, pers. obs.).

**Figure 1.**
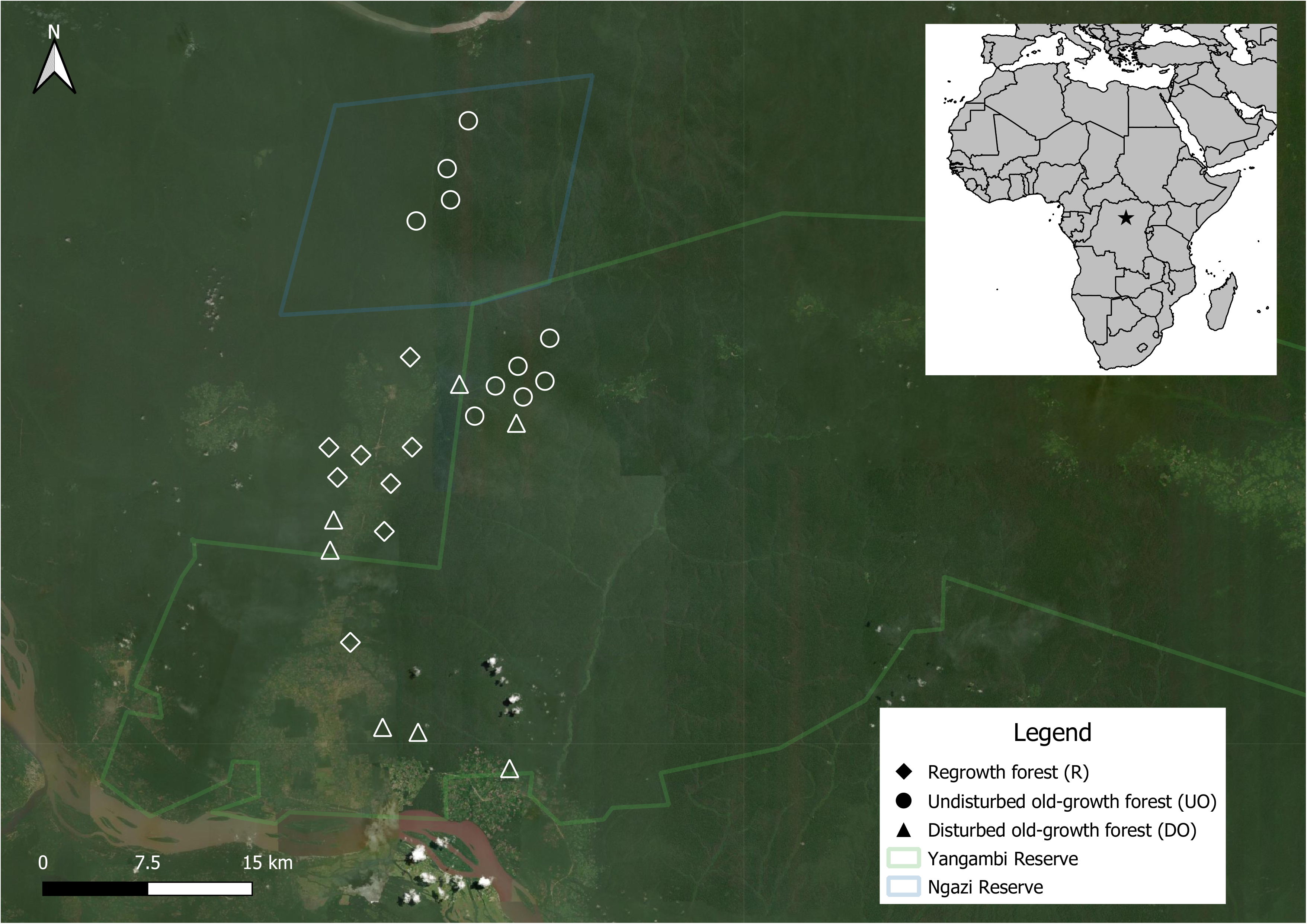
Location of the 25 forest inventory plots across the three forest categories in the Yangambi region (DR Congo). Base layer: ESRI SAT.

### 2.2. Vegetation survey and plant trait quantification

Five quadrats (25 m × 25 m) were randomly established in each forest inventory plot, resulting in a total of 125 sampling quadrats. In each quadrat, the abundance and diameter at breast height (DBH) of all woody species with a DBH ≥ 5 cm were measured. The quadrats were geolocated (Garmin GPSMAP 64 series) and crown closure (%) was quantified from four readings in the cardinal directions with a model A convex spherical densiometer.

The specific leaf area (SLA) of all taxa identified at species level was assessed on dried herbarium samples that were collected during the vegetation surveys. For species without samples collected in the field, dry Central-African herbarium specimens from the Meise Botanic Garden herbarium were measured. The measurements were performed following Pérez-Harguindeguy et al. (2016). Leaf area was assessed using leaf disks (4.9 cm^2^), to minimise damage to herbarium specimens, or otherwise by a whole leaf scan using ImageJ (Rueden et al., 2017). The obtained disks and whole leaves were weighted afterwards. Per species five replicates were used. In addition to the SLA, the wood density (WD) of 147 species was compiled from the literature (Table S3) (Carsan et al., 2012; Harja et al., 2021; RMCA, 2021).

### 2.3. Data analyses

Statistical analyses were performed using R software (R Core Team, 2020). Based on the five quadrats, for each plot, the number of trees, species richness (Hill’s N_0_), species diversity (Hill’s N_1_ or the exponential of the Shannon entropy) and evenness (Hill’s N_1_/log(Hill’s N_0_)) were computed. The estimates were standardised for sample completeness, as sample sizes differed between plots. Sample completeness represents the proportion of the total number of individuals in a unit that belong to the species in the sample, and contrary to sample size standardisation, this standardisation method allows less biased comparisons (Chao et al., 2012). Extrapolation of the estimates was done to twice the reference sample size per plot following Chao & Jost (2012). Calculations were based on the collected species abundance data with 9999 bootstraps using the *iNEXT* function of the iNEXT package (Hsieh et al., 2016). Subsequently, the different metrics and the crown closure were tested for differences between the three forest categories using Kruskal-Wallis rank sum tests and pairwise Wilcoxon rank sums tests with correction for multiple testing, as the assumptions for parametric testing were not met.

Dissimilarity of the tree community composition across the forest categories was visualised using a non-metric multidimensional scaling (NMDS) based on the Bray-Curtis distance matrix, the most suitable distance for abundance data (Ricotta et al., 2017). A two-axis solution provided a stress level (0.13) well below the 0.20 threshold. The ordination was performed with the *metaMDS* function in the vegan package (Oksanen et al., 2020). Subsequently, a permutational multivariate analysis of variance with 9999 permutations was performed to test for differences between the species composition of the three different forest categories using the *adonis* function in the Vegan package (Oksanen et al., 2020). Pairwise comparisons of the different forest groups with 9999 permutations and Bonferroni correction were performed with the *pairwise.adonis.function* in the pairwiseAdonis package (Arbizu, 2017). Tree species indicative for each of the three forest categories were identified based on their indicator value (Dufrene et al., 1997). This value was calculated using the *multipatt* function in the indicspecies package with 9999 permutations.

Community-weighted means (CWM) for each plot were calculated for the SLA and the WD. The community-weighted means are known to be very responsive to environmental change and highly sensitive to disturbance (Carreño-Rocabado et al., 2012). Both variables met the assumptions for parametric testing (normality > 0.05 & homogeneity > 0.05). To evaluate differences in community-weighted mean SLA and WD among forest categories, a one-way analysis of variance (ANOVA) was conducted. Differences were further investigated using Tukey post hoc tests.

Forest structure was evaluated using the measured DBHs. Trees were categorised in ten size classes according to their DBH. An interval of 10 cm was used in the first nine classes, starting from 5 cm DBH. The tenth class had a wider interval, aggregating individuals with a DBH over 95 cm. Subsequently, diameter frequency distribution was compared among forest categories with pearson’s chi-squared tests with simulated p-values based on 9999 replicates using the *chisq.test* function in the stats package. The chi-square test was preferred over the frequently used Kolmogorov-Smirnov (Mitchell, 1971). The diameter frequency distribution comparison was complemented by the calculation of the structural diversity, which is a single measure summarising the diameter distribution (McElhinny et al., 2005). Structural diversity was quantified by the first order Hill number which was standardised for sample completeness following the same methodology as outlined above. Calculation and extrapolation of structural diversity was done with the *iNEXT* function with 9999 bootstraps in the iNEXT package, as above.

## 3. RESULTS

### 3.1. Species richness and diversity

A total of 7375 woody plant individuals (DBH ≥ 5 cm) from 211 species was recorded across the 25 forest inventory plots. Of the 211 species, 79.15% was identified to species level, 14.22% only to genus level and 6.63% were unidentified species. Crown closure (*χ*^2^ = 2.21, *P* > 0.05) and hill’s N_0_ (*χ*^2^ = 1.98, *P* > 0.05) were not significantly different among the forest categories. Whereas tree abundance (*χ*^2^ = 10.90, *P*< 0.01), Hill’s N_1_ (*χ*^2^ = 9.78, *P* < 0.01) and evenness (*χ*^2^ = 10.06, *P* < 0.01) did differ between the forest categories (Figure 2). Tree abundance was 17.2% lower (*P* < 0.05) in DO forests and 24.3% (*P* < 0.05) lower in R forests, when compared to UO forests. Species diversity and evenness were 33.6% (*P* < 0.001) and 31.9% (*P* < 0.001) higher, respectively, in DO forests as compared to UO forests.

**Figure 2.**
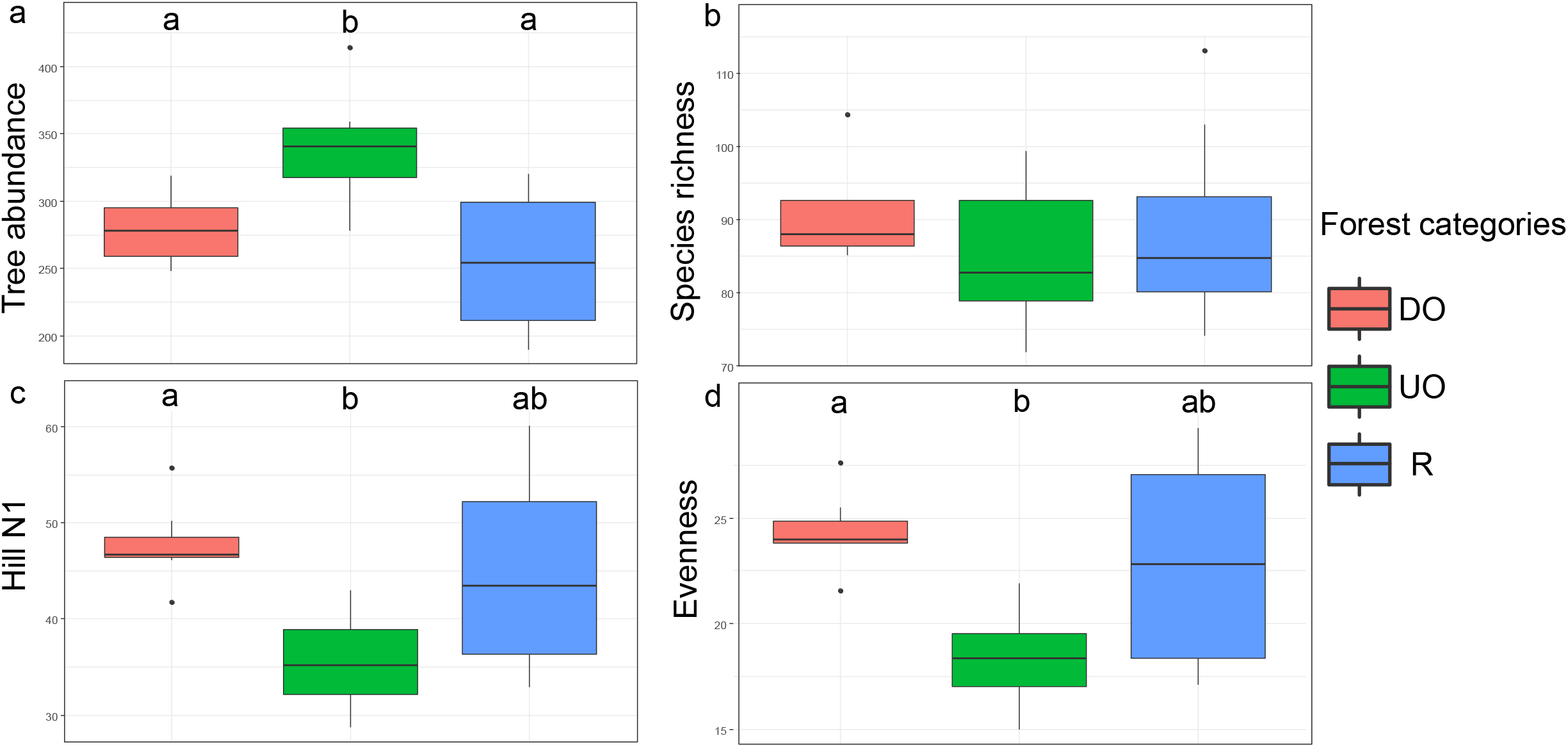
Boxplots comparing tree abundance and tree species diversity metrics across old-growth undisturbed forest (OU), old-growth disturbed forest (OD), and regrowth forest (R) in the Yangambi region (DR Congo). Hinges represent the 25th, 50th and 75th percentiles, respectively. Whiskers extend to maximum 1.5 times the interquartile range. Letters code for significant differences between groups. (a) tree abundance, (b) species evenness, (c) species richness, (d) Hill’s N1.

### 3.2. Species composition

The NMDS ordination of the 25 forest inventory plots showed a clear separation of the UO forests from the other forest categories, whereas also the R forests tended to cluster at the outsides of the graph (Figure 3). This was supported by the permutational multivariate analysis of variance which indicated differences between categories (*R*^*2*^ = 0.253; *P* < 0.001). Specifically, species composition significantly differed between DO forests and UO forests (*R*^*2*^= 0.207; *P* < 0.001), and between R forests and UO forests (*R*^*2*^ = 0.267; *P* < 0.001). Across the 25 forest inventory plots, 27 of the 167 to species level identified trees were indicative based on their indicator value (both *P* < 0.05 and indicator value > 0.20). Of these indicative species, 17 were indicative for UO forest (e.g. *Cola griseiflora*), 4 for DO forest (e.g. *Leplaea cedrata*) and 6 for R forest (e.g. *Heisteria parvifolia*) (Table S2).

**Figure 3.**
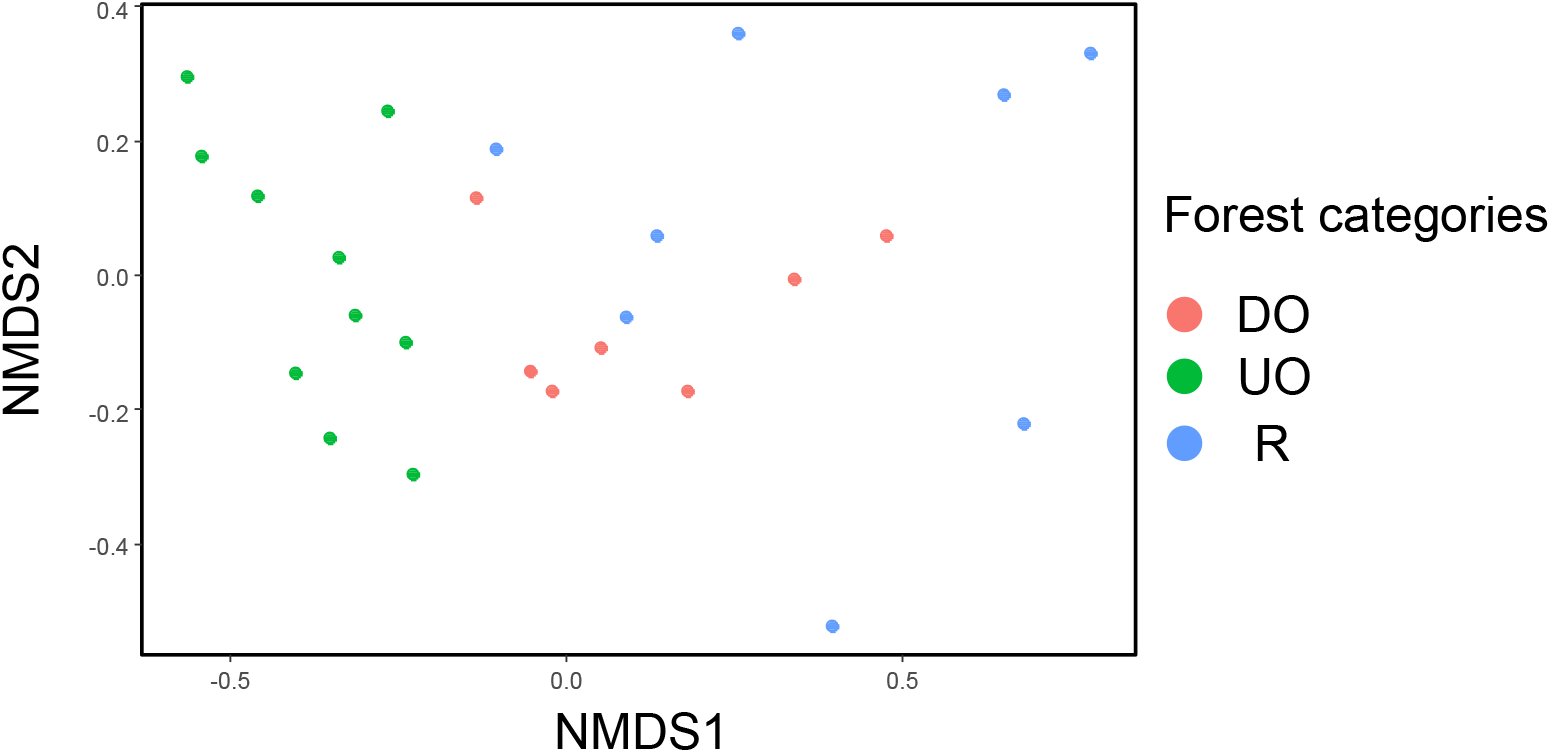
Non-metric multidimensional scaling (NMDS) of 25 forest inventory plots in the Yangambi region (DR Congo) across the three forest categories: undisturbed old-growth forest (UO), disturbed old-growth forest (DO), and regrowth forest (R). Ordination is based on the Bray-Curtis distance matrix calculated from the tree species abundances and represents the woody species (DBH ≥ 5 cm) community composition.

### 3.3. Specific leaf area and wood density

Whereas the community-weighted mean for SLA did not differ between forest categories (*F*= 1.38, *P*> 0.05), community-weighted mean WD did (*F*= 9.03, *P* < 0.01) (Figure 4). In DO forests, community-weighted mean WD was 5.0 % lower (*P*<0.05) and in R forests it was 6.2 % lower (*P*< 0.01) as compared to UO forests.

**Figure 4.**
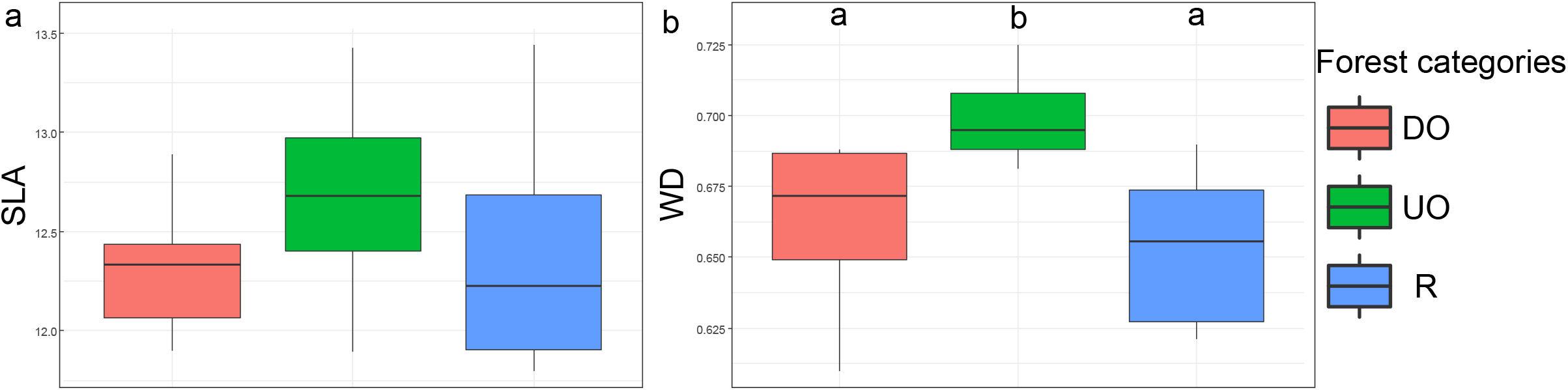
Boxplots comparing the community weighted means of specific leaf area (SLA) and wood density (WD) across the three forest categories. Hinges represent the 25^th^, 50^th^ and 75^th^ percentiles, respectively. Whiskers extend to maximum 1.5 times the interquartile range. Letters code for significant differences between groups.

### 3.4. Forest structure

Based on the diameter distribution there were more small sized trees in UO forests than in DO forests (*χ*^2^= 34.51; *P* < 0.001) and in R forests (*χ*^2^=61.54; *P*< 0.001) (Figure 5). These differences between categories were also present for the structural diversity, here represented by Hill’s N_1_, (*χ*^2^ = 13.76; *P* < 0.01), where UO forests differed from DO forests (*P* < 0.01) and from R forests (P < 0.01).

**Figure 5.**
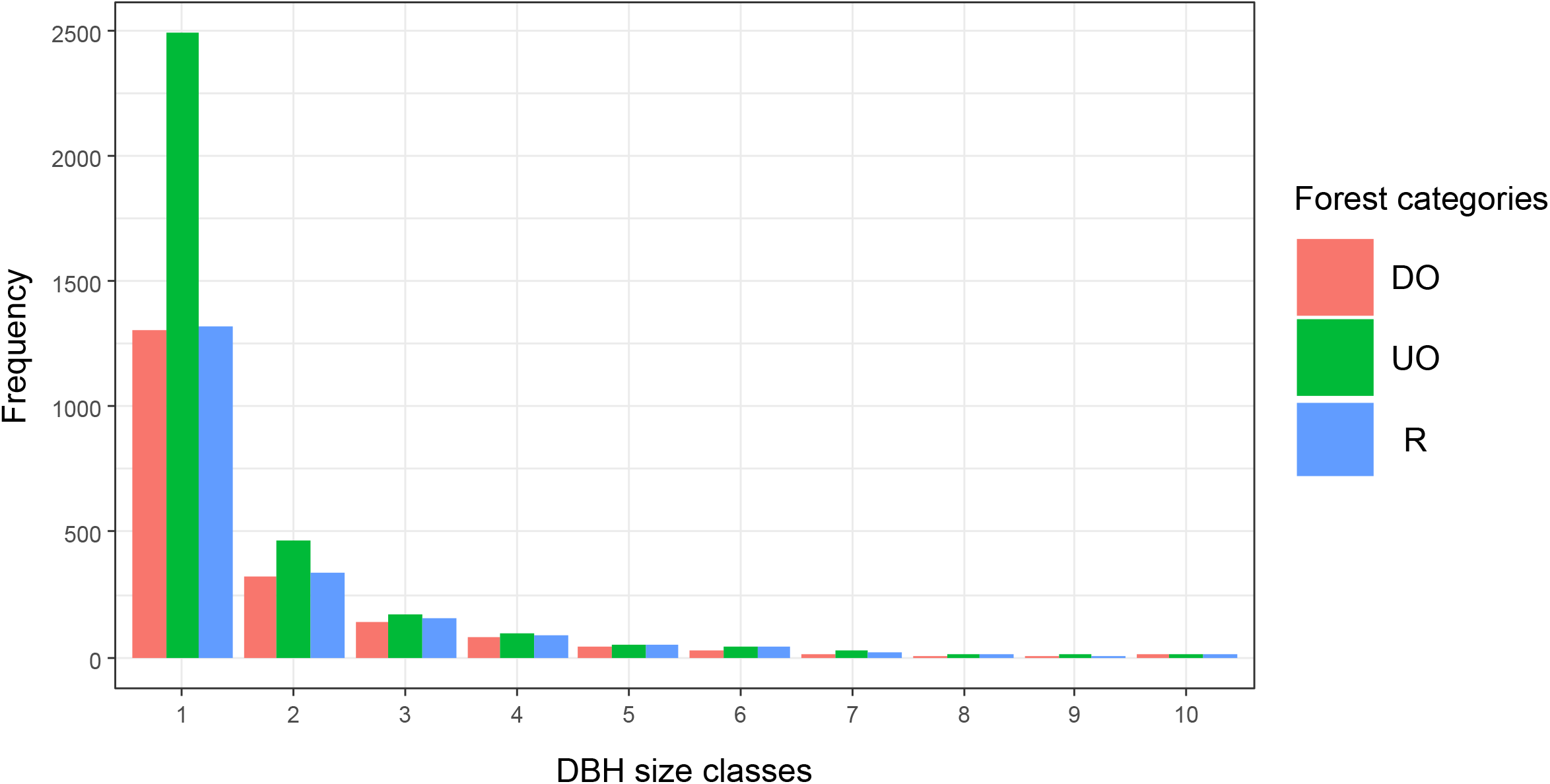
Observed frequencies among the ten DBH size categories compared among the three forest categories. Size classes one to nine have a 10 cm interval, while the tenth has a wider interval, grouping individuals with a DBH of over 95 cm.

## 4. DISCUSSION

Understanding how anthropogenic disturbance affects forest structure, species richness and species composition is of major importance for biodiversity conservation in globally underexplored biodiversity hotspots such as the Congo Basin. Based on ground surveys and historical land-use maps, we provided insights in the effects of forest degradation and of forest regeneration following agricultural abandonment in the Yangambi area. We found significant effects of human disturbance on tree species composition and tree abundance, and provided evidence that 60 years after agricultural abandonment, the overall tree species composition is still different from that of undisturbed old-growth forest.

### 4.1. Forest disturbance

Whereas the crown closure did not reflect anthropogenic induced changes in the study area, our ground survey showed that human activities clearly cause shifts in community composition and forest structure, even though, the number of woody species was not affected, and species diversity even increased following disturbance. The latter likely indicates that the species abundance distribution is less skewed in disturbed forests. Indeed, the evenness index showed that the species abundance is more evenly distributed in disturbed forest than in undisturbed forest. Combined with the observation that disturbed forests contained significantly less tree individuals, these findings suggest that selective logging, in which exploitable species are selectively removed, is the main cause behind these patterns. For example, *Prioria oxyphylla* and *Scorodophloeus zenkeri* were both much less abundant in disturbed forests and are extensively used as construction material. Moreover, in DR Congo the latter is also exploited for its medicinal properties and is used to make charcoal (Plant Resources of Tropical Africa, 2021). Furthermore, in our study area, species such as *Dialium pachyphyllum* and *Chlamydocola chlamydantha* were more abundant in undisturbed old-growth forest and are indicators of old-growth forest (Meunier et al., 2015).

Disturbed old-growth forest on the other hand, was characterised by shadow-tolerant tree species of which the wood is non-exploitable. For example, species of the genus *Cola* were found in both disturbed and undisturbed old-growth forest. These species are not selectively logged because their wood is of little use due to a low DBH and highly irregular trunks. Rather, these species are conserved, especially *Cola acuminata*, because of their highly valued nuts which have stimulating and medicinal properties (Meunier et al., 2015). Likewise, the abundance of *Diospyros* spp. is indicative for old growth forests and are non-exploitable, except *Diospyros crassiflora* of which the wood is highly valued (Meunier et al., 2015). Similarly, studies in other tropical regions have observed changes in dominating species following disturbance (Baboo et al., 2017), with a decrease in old-growth characteristic species (Makana et al., 2006) and an increase in secondary forest species (Ifo et al., 2016).

Logging in tropical Africa is generally limited to one or two trees per hectare (Bousfield et al., 2020) and is recognised to be the primary driver of forest degradation in Africa (Hosonuma et al., 2012; Jayathilake et al., 2021). However, its influence on species diversity is still unclear. A study in Congolese forests comparable to those in Yangambi, compared disturbed and undisturbed forests in two distinct locations (Makana et al., 2006). Whereas a significant influence of disturbance was detected in one location, similar values for species richness and species diversity were detected in the second location. Likewise, a clear effect of disturbance on species diversity was detected in tropical forests in the Republic of the Congo (Ifo et al., 2016) and India (Baboo et al., 2017), but not in a tropical forest in Bolivia (Carreño-Rocabado et al., 2012). Given that the response of species diversity to disturbance is highly variable, solely assessing the impact of anthropogenic disturbance on tropical vegetation through species diversity can mask its severity (Edwards et al., 2014). Our results confirm that the tree species composition in Yangambi forests was significantly affected by anthropogenic disturbance. In contrast to species diversity, species composition is an excellent indicator of forest degradation (Imai et al., 2014). Whereas many studies across different tropical regions have reported varying responses of species diversity to forest disturbance, they all reported significant shifts in the species composition following anthropogenic disturbance (Baboo et al., 2017; Carreño-Rocabado et al., 2012; Ifo et al., 2016; Makana et al., 2006). These disturbances may change tree community composition to such a degree that species interactions are weakened, although not fully lost (Poulsen et al., 2013).

Ecosystem functioning is not only mediated by species diversity, but also depends on species’ functional traits (de Bello et al., 2021). Hence, assessing the effect of disturbance on community-weighted mean functional traits can provide more ecologically relevant information than only using taxonomic data (Katovai et al., 2012). We expected community-weighted mean SLA to increase and WD to decrease following disturbance (Rocha-Santos et al., 2019; Tscharntke et al., 2012). Remarkably, no significant differences were observed in community-weighted mean SLA between forest categories. A higher SLA would imply enhanced foraging to maximally capture light resources (Poorter & Bongers, 2006). In our study, the different forest categories did not differ in light availability as the canopy remained untouched. It has indeed been established by Carreño-Rocabado et al. (2012) that changes in community-weighted mean functional traits in tropical forests are mainly driven by increased light availability. Furthermore, although disturbed and undisturbed old-growth forests were characterised by different indicator species assemblages, the majority can still be classified as shade-tolerant, which is commonly associated with low SLA (Poorter, 2009). In contrast to community-weighted mean SLA, community-weighted mean WD did significantly differ between disturbed and undisturbed old-growth forest, suggesting that trees with high WD of high construction quality, have been selectively removed from the disturbed forests (Saranpää, 2003).

Like the majority of African tropical forests, the mean diameter distribution in the forests in the Yangambi region, could be described by the reverse J-shaped model (Fayolle et al., 2014). Our results indicate that anthropogenic disturbance significantly influences this reverse J-shaped tree size distribution. Particularly, the reverse J-shape was more pronounced in undisturbed old-growth forests than in disturbed old-growth forests which contained significantly less trees in the first size class. It has been recognised that the shape of the tree size distribution can provide important information on the status of tree regeneration, and in which the reverse J-shape is an indicator for a substantial regeneration (Gebeyehu et al., 2019). Similar results of reduced forest regeneration were obtained in other tropical regions (Baboo et al., 2017; Hall et al., 2003; Makana et al., 2006). The influence of anthropogenic disturbance on tree regeneration is three-fold. Firstly, selective logging results in the reduction of the overall number of trees and, accordingly, in the reduction of seed sources (Makana et al., 2006; Medjibe et al., 2014). Combined with the fact that pollination and seed dispersal are severely affected by disturbance (Neuschulz et al., 2016), this may result in a decreased forest regeneration. Secondly, selective logging comes with increased trampling, felling damage, and soil compaction affecting seedling establishment and survival (Baboo et al., 2017; Chapman et al., 1997). Lastly, disturbance alters the species composition, possibly enhancing the number of poorly regenerating species (Baboo et al., 2017) and changing species interactions (e.g. competition), ultimately jeopardising forest regeneration.

### 4.2. Forest regeneration following agricultural abandonment

Based on historical land-use maps we were able to assess the recovery of abandoned agricultural fields and observed that regrowth forests recovered in terms of species diversity to levels comparable to those found in disturbed old-growth forests, and even to those in undisturbed old-growth forests. They only differed from the latter in terms of the number of tree individuals. These findings on the recovery of species diversity are in line with other studies in the Congo Basin (Bauters et al., 2019; Makelele et al., 2021) and in other tropical regions (Letcher et al., 2009; Mahayani et al., 2020). In contrast to species diversity, we found that the species composition in regrowth forest was still significantly different from that in undisturbed old-growth forest. Similar to our results, the species composition of young regrowth forest (25-30 years) differed from old-growth forest in the Maringa-Lopori-Wamba forest landscape in DR Congo, whereas old regrowth forests (150-300 years) were very close to old-growth forests (Bauters et al., 2019). Likewise, a recent study in the Yoko forest reserve, at approximately 30 km Southeast of Kisangani, showed that the species composition of 60-year-old regrowth forests was still significantly different from local old-growth forests (Makelele et al., 2021). Our results are furthermore in line with studies in the Neotropics (Oberleitner et al., 2021) and tropical Asia (Hayward et al., 2021).

Despite having a similar species composition as disturbed old-growth forests, regrowth forests were characterised by a different indicator species assemblage, including for example *Myrianthus arboreus*, a late pioneer species (Plant Resources of Tropical Africa, 2021). Additionally, *Combretum lokele* and *Dacryodes edulis* were also characteristic for regrowth forests. Both species are commonly found in villages and are indicative for the anthropogenic influence in these forests (Meunier et al., 2015).

The forest structure in the regrowth forests was comparable to disturbed old-growth forest, yet still significantly different from undisturbed old-growth forest. These differences were not caused by the lower abundance of large-sized trees in regrowth forests, as would be expected because these are slower to recover (Bauters et al., 2019; Makelele et al., 2021). On the contrary, the observed differences can be explained by the higher frequency of small-sized trees in undisturbed old-growth forests. This implies that, like in the disturbed old-growth forests, tree regeneration in regrowth forests is hampered. Similar to other tropical regions (Bauters et al., 2019; Makelele et al., 2021), forests in the Yangambi landscape seem to initially recover rapidly to a close canopy forest following agricultural abandonment, but further forest succession may be hampered by ongoing anthropogenic disturbance. Overall, anthropogenic disturbance not only causes the degradation of existing forests, but also jeopardises their recovery.

### 4.3. Implications

We found that the tree species composition and forest structure in disturbed forest were significantly different from undisturbed old-growth forest. Likewise, in a little more than 60 years following agricultural abandonment, species diversity was able to recover, but species composition and forest structure remained significantly different in these regrowth forests. Additionally, our data suggest that disturbed old-growth forest and regrowth forest are as productive as undisturbed old-growth forest, but are not able to store as much carbon, based on the SLA and WD, respectively (Chave et al., 2009; Madani et al., 2018). Yet, although primary forests are deemed irreplaceable (Gibson et al., 2011), studies show that such regrowth forests can play a crucial role in maintaining biodiversity and ecosystem services (Crouzeilles et al., 2016; Van de Perre et al., 2018). In short, our study highlights the need of proper conservation, especially of remaining old-growth forests, but also of regrowth forests in Yangambi. Although the studied forests are part of the Yangambi Man and Biosphere reserve, this provides little protection from forest degradation (Koy et al., 2019). A global analysis has pointed out that for protected areas to be effective, they should be coupled with increased law enforcement, increased efforts to reduce corruption and increased stakeholder engagement (Geldmann et al., 2019). We argue that human pressure on Yangambi’s biodiversity can be reduced by providing alternatives for bushmeat and wood fuel which will inevitably reduce hunting and selective logging, respectively, and thus forest degradation.

## ACKNOWLEDGMENTS

We would like to thank the Ministère de L’Environnement et Développement Durable (MEDD) and Institut National pour l’Étude et la Recherche Agronomiques (INERA) and the FORETS project, which is financed by the European Union, for their help in the field. We would also like to thank both Hans Beeckman and Tom De Mil of the Royal Museum for Central Africa (RMCA) for their help in retrieving wood density data. This study was funded by Research Foundation-Flanders, research mandate granted to JD (FWO; 1125221N) and research project granted to OH (FWO; G090719N), and the Foundation for the promotion of biodiversity research in Africa, granted to JD and YH (SBBOA, www.sbboa.be).

## CONFLICT OF INTEREST

All authors confirm that there is no conflict of interest regarding the publication of this article.

## AUTHOR CONTRIBUTIONS

OH, FV and JD designed this study. JD, JA, RM, YH and JK participated in fieldwork. JD analysed the data. FV, OH and JD wrote the manuscript. All authors contributed to finalising the manuscript.

## DATA AVAILABILITY STATEMENT

The data that support the findings of this study will be made openly available in Zenodo

## SUPPORTING INFORMATION

Additional supporting information can be found online in the Supporting Information section at the end of the article.

APPENDIX S1 Plot level data

APPENDIX S2 Indicator species per forest category

APPENDIX S3 Plant trait values per species

## Notes

### Competing Interest Statement

The authors have declared no competing interest.

