## Supplementary info for "Effects of forest disturbance and regeneration on tree species composition and traits in DR Congo"

Appendix S1 plot level data. Cat: forest category including undisturbed old-growth forest (UO), disturbed old-growth forest (DO), and regrowth forest (R); Plot: forest inventory plot; m: sample size; sc: sample completeness; q0: hill number of zeroth order; q1: hill number of first order; EV: evenness; NT: number of trees; OV: overstory density; WD: community-weighted mean of wood density; SLA: community-weighted mean of specific leaf area; SD: structural diversity

| **Cat** | **Plot** | **m** | **sc** | **q0** | **q1** | **EV** | **NT** | **OV** | **WD** | **SLA** | **SD** |
| --- | --- | --- | --- | --- | --- | --- | --- | --- | --- | --- | --- |
| DO | 1 | 592 | 0.96 | 92 | 46.81 | 23.81 | 296 | 89.08 | 0.686 | 11.898 | 2.82 |
| DO | 2 | 588 | 0.96 | 86 | 46.10 | 23.80 | 294 | 92.72 | 0.688 | 11.924 | 2.93 |
| DO | 3 | 556 | 0.95 | 88 | 46.64 | 23.98 | 278 | 91.37 | 0.672 | 12.890 | 2.92 |
| R | 4 | 444 | 0.95 | 74 | 38.19 | 20.42 | 222 | 90.80 | 0.660 | 13.111 | 4.04 |
| DO | 5 | 514 | 0.96 | 85 | 46.73 | 24.21 | 257 | 90.28 | 0.654 | 12.209 | 3.17 |
| R | 6 | 572 | 0.97 | 90 | 52.72 | 26.99 | 286 | 91.68 | 0.690 | 11.840 | 3.19 |
| R | 7 | 622 | 0.99 | 83 | 32.96 | 17.14 | 209 | 73.63 | 0.621 | 12.539 | 3.25 |
| R | 8 | 418 | 0.94 | 82 | 52.05 | 27.22 | 311 | 90.54 | 0.626 | 11.925 | 2.95 |
| R | 9 | 380 | 0.92 | 86 | 48.81 | 25.25 | 190 | 90.33 | 0.652 | 13.441 | 3.89 |
| R | 10 | 640 | 0.96 | 103 | 36.97 | 18.37 | 320 | 91.21 | 0.628 | 11.930 | 2.92 |
| DO | 11 | 496 | 0.96 | 86 | 41.77 | 21.57 | 248 | 93.34 | 0.610 | 12.335 | 3.25 |
| DO | 12 | 638 | 0.97 | 93 | 50.18 | 25.51 | 319 | 93.45 | 0.688 | 12.480 | 3.12 |
| R | 13 | 424 | 0.94 | 75 | 34.46 | 18.35 | 212 | 89.86 | 0.683 | 12.520 | 2.80 |
| R | 14 | 590 | 0.93 | 113 | 60.06 | 29.24 | 295 | 94.28 | 0.671 | 11.797 | 3.98 |
| DO | 15 | 522 | 0.95 | 104 | 55.76 | 27.62 | 261 | 91.89 | 0.644 | 12.390 | 3.12 |
| UO | 16 | 712 | 0.98 | 78 | 31.59 | 16.70 | 356 | 90.02 | 0.710 | 12.794 | 3.29 |
| UO | 17 | 700 | 0.98 | 83 | 35.72 | 18.61 | 350 | 89.91 | 0.710 | 12.952 | 2.44 |
| UO | 18 | 828 | 0.98 | 82 | 28.79 | 15.03 | 414 | 92.51 | 0.701 | 13.424 | 2.62 |
| UO | 19 | 718 | 0.98 | 75 | 33.94 | 18.07 | 359 | 91.73 | 0.725 | 11.893 | 2.34 |
| UO | 20 | 556 | 0.95 | 96 | 37.81 | 19.08 | 278 | 91.84 | 0.700 | 12.106 | 3.13 |
| UO | 21 | 622 | 0.97 | 82 | 34.69 | 18.13 | 311 | 91.21 | 0.681 | 12.521 | 2.84 |
| UO | 22 | 674 | 0.97 | 92 | 43.01 | 21.91 | 337 | 91.47 | 0.688 | 12.564 | 2.56 |
| UO | 23 | 580 | 0.95 | 93 | 41.68 | 21.17 | 290 | 90.59 | 0.690 | 12.361 | 2.65 |
| UO | 24 | 688 | 0.99 | 72 | 30.40 | 16.37 | 344 | 95.32 | 0.683 | 13.244 | 2.56 |
| UO | 25 | 676 | 0.95 | 99 | 39.31 | 19.68 | 338 | 94.28 | 0.689 | 12.980 | 2.59 |

Appendix S2 Indicator species across the three forest categories including undisturbed old-growth forest (UO), disturbed old-growth forest (DO), and regrowth forest (R), with their corresponding indicator value and p-value.

| SPECIES | CAT | VALUE | P |
| --- | --- | --- | --- |
| *Leplaea cedrata* | DO | 0.711 | 0.0394 |
| *Cola acuminata* | DO | 0.697 | 0.0134 |
| *Diospyros iturensis* | DO | 0.697 | 0.0137 |
| *Diospyros crassiflora* | DO | 0.690 | 0.0128 |
| *Cola griseiflora* | UO | 0.831 | 0.0001 |
| *Cola gigantea* | UO | 0.823 | 0.0016 |
| *Drypetes morocarpa* | UO | 0.792 | 0.0023 |
| *Chlamydocola chlamydantha* | UO | 0.785 | 0.0058 |
| *Drypetes gossweileri* | UO | 0.779 | 0.0015 |
| *Garcinia punctata* | UO | 0.759 | 0.0082 |
| *Synsepalum subcordatum* | UO | 0.756 | 0.0078 |
| *Dialium pachyphyllum* | UO | 0.751 | 0.0096 |
| *Cynometra hankei* | UO | 0.730 | 0.0454 |
| *Prioria oxyphylla* | UO | 0.721 | 0.0174 |
| *Monodora myristica* | UO | 0.713 | 0.0449 |
| *Greenwayodendron suaveolens* | UO | 0.711 | 0.0143 |
| *Massularia acuminata* | UO | 0.701 | 0.0199 |
| *Scorodophloeus zenkeri* | UO | 0.695 | 0.0368 |
| *Xylopia phloiodora* | UO | 0.665 | 0.0476 |
| *Cola urceolata* | UO | 0.632 | 0.0280 |
| *Tessmannia anomala* | UO | 0.632 | 0.0237 |
| *Heisteria parvifolia* | R | 0.866 | 0.0003 |
| *Myrianthus arboreus* | R | 0.814 | 0.0018 |
| *Carapa procera* | R | 0.760 | 0.0318 |
| *Turraeanthus africanus* | R | 0.728 | 0.0235 |
| *Combretum lokele* | R | 0.707 | 0.0067 |
| *Dacryodes edulis* | R | 0.681 | 0.0376 |

Appendix S3 Trait values per species. WD: Wood Density; SLA: Specific Leaf Area; HB ref: herbarium reference; WD ref: source of Wood Density data. Species names following the African Plant Database (Conservatoire et Jardin botaniques de la Ville de Genève and South African National Biodiversity Institute, 2021).

| Species | WD | SLA | HB ref | WD ref |
| --- | --- | --- | --- | --- |
| *Afrostyrax lepidophyllus* | 0.722 | 29.454 | 0 000005 60437 1 | RMCA (DRYAD, 2021) |
| *Afzelia bipindensis* | 0.615 | 10.983 | BR0000015254221V | RMCA (DRYAD, 2021) |
| *Aidia micrantha* | 0.931 | 11.665 | BR0000013018009 | RMCA (DRYAD, 2021) |
| *Albizia adianthifolia* | 0.631 | 7.682 | BR0000015842749 | RMCA (DRYAD, 2021) |
| *Albizia ferruginea* | 0.608 | 12.563 | BR0000017659031 | RMCA (DRYAD, 2021) |
| *Albizia gummifera* | 0.612 | 20.396 | BR0000016024120 | RMCA (DRYAD, 2021) |
| *Albizia zygia* | 0.512 | 11.454 | BR0000016041707 | RMCA (DRYAD, 2021) |
| *Allanblackia floribunda* | 0.681 | 11.112 | BR0000016603455 | Harja et al. (2021) |
| *Alstonia boonei* | 0.408 | 8.206 | BR00000147045154 | RMCA (DRYAD, 2021) |
| *Amphimas pterocarpoides* | 0.851 | 23.662 | BR0000016720497 | RMCA (DRYAD, 2021) |
| *Angylocalyx pynaertii* | 0.67 | 9.491 | BR0000016706231 | RMCA (DRYAD, 2021) |
| *Anonidium mannii* | 0.427 | 14.37 | BR000001514159 | RMCA (DRYAD, 2021) |
| *Anopyxis klaineana* | 0.81 | 6.314 | BR0000017685429 | RMCA (DRYAD, 2021) |
| *Anthonotha macrophylla* | 0.59 | 11.169 | BR0000016133709 | RMCA (DRYAD, 2021) |
| *Antiaris toxicaria* | 0.443 | 6.112 | BR0000016843929 | RMCA (DRYAD, 2021) |
| *Aoranthe cladantha* | 0.765 | 15.422 | BR0000019099682 | RMCA (DRYAD, 2021) |
| *Aphanocalyx microphyllus* | NA | 5.767 | BR0000016129900V | NA |
| *Barteria fistulosa* | 0.673 | 12.071 | BR0000017638104 | RMCA (DRYAD, 2021) |
| *Blighia unijugata* | 0.684 | 13.179 | BR0000018152142 | RMCA (DRYAD, 2021) |
| *Blighia welwitschii* | 0.745 | 9.649 | BR0000013258551 | RMCA (DRYAD, 2021) |
| *Bridelia micrantha* | NA | 12.015 | BR0000015812896 | NA |
| *Canarium schweinfurthii* | 0.468 | 9.304 | BR0000014228261 | RMCA (DRYAD, 2021) |
| *Carapa procera* | 0.721 | 6.312 | BR0000013595647 | RMCA (DRYAD, 2021) |
| *Celtis mildbraedii* | 0.74 | 8.548 | BR0000013285821 | RMCA (DRYAD, 2021) |
| *Celtis tessmannii* | 0.659 | 9.454 | BR0000020351632 | RMCA (DRYAD, 2021) |
| *Chaetocarpus africanus* | 0.811 | 11.675 | BR0000015794390 | RMCA (DRYAD, 2021) |
| *Chlamydocola chlamydantha* | NA | 10.168 | BR0000018733488 | NA |
| *Chrysophyllum africanum* | 0.61 | 10.146 | BR0000013987022 | Harja et al. (2021) |
| *Chrysophyllum lacourtianum* | 0.7 | 8.758 | BR0000020188405 | Carsan et al. (2012) |
| *Chrysophyllum pruniforme* | 0.64 | 11.308 | BR0000020189204 | Carsan et al. (2012) |
| *Chytranthus carneus* | 0.717 | 14.191 | BR0000018160604 | RMCA (DRYAD, 2021) |
| *Chytranthus macrobotrys* | 0.691 | 16.061 | BR0000018161908 | RMCA (DRYAD, 2021) |
| *Cleistanthus caudatus* | 0.896 | 18.741 | BR0000015247087V | Harja et al. (2021) |
| *Coelocaryon botryoides* | NA | 11.644 | BR0000013439408 | NA |
| *Coelocaryon preussii* | 0.468 | 10.559 | BR0000013966669 | RMCA (DRYAD, 2021) |
| *Coffea canephora* | 0.71 | 25.436 | JDP0230 | RMCA (DRYAD, 2021) |
| *Cola acuminata* | 0.64 | 9.335 | BR0000019774275 | RMCA (DRYAD, 2021) |
| *Cola altissima* | NA | 11.709 | BR0000019680255 | NA |
| *Cola bruneelii* | NA | 17.841 | BR0000013672645 | NA |
| *Cola gigantea* | 0.59 | 6.568 | BR0000020100391 | RMCA (DRYAD, 2021) |
| *Cola griseiflora* | 0.689 | 13.897 | BR0000013631611 | RMCA (DRYAD, 2021) |
| *Cola lateritia* | 0.609 | 9.627 | BR0000019981123 | RMCA (DRYAD, 2021) |
| *Cola marsupium* | NA | 19.058 | BR0000020015916 | NA |
| *Cola urceolata* | NA | 8.899 | BR0000019982199 | NA |
| *Combretum lokele* | 0.717 | 9.116 | BR0000013692183 | RMCA (DRYAD, 2021) |
| *Copaifera mildbraedii* | 0.709 | 7.57 | 0 000006 87232 | RMCA (DRYAD, 2021) |
| *Cynometra hankei* | 0.763 | 20.197 | BR0000013624869 | RMCA (DRYAD, 2021) |
| *Cynometra pedicellata* | NA | 7.326 | BR0000016551541 | NA |
| *Dacryodes edulis* | 0.532 | 15.576 | BR0000014203947 | RMCA (DRYAD, 2021) |
| *Desplatsia chrysochlamys* | 0.426 | 24.475 | BR0000019963600 | RMCA (DRYAD, 2021) |
| *Desplatsia dewevrei* | 0.395 | 21.656 | BR0000020059361 | RMCA (DRYAD, 2021) |
| *Dialium pachyphyllum* | 0.923 | 8.049 | BR0000013637620 | Harja et al. |
| *Dichostemma glaucescens* | 0.6 | 8.404 | BR0000016228986 | RMCA (DRYAD, 2021) |
| *Diogoa zenkeri* | 0.715 | 7.302 | BR0000017035712 | RMCA (DRYAD, 2021) |
| *Diospyros crassiflora* | 0.716 | 9.854 | BR0000013251338 | RMCA (DRYAD, 2021) |
| *Diospyros hoyleana* | 0.717 | 9.897 | BR0000015608345 | RMCA (DRYAD, 2021) |
| *Diospyros iturensis* | 0.785 | 11.789 | BR0000013300692 | RMCA (DRYAD, 2021) |
| *Dracaena arborea* | NA | 13.189 | BR0000013486368 | NA |
| *Drypetes cinnabarina* | 0.877 | 16.98 | BR0000015261724 | RMCA (DRYAD, 2021) |
| *Drypetes gossweileri* | 0.744 | 11.498 | BR0000013609733 | RMCA (DRYAD, 2021) |
| *Drypetes ituriensis* | NA | 12.195 | BR0000015790019 | NA |
| *Drypetes morocarpa* | NA | 12.627 | 0 000006 77124 9 | NA |
| *Entandrophragma angolense* | 0.579 | 7.393 | BR0000013596316 | RMCA (DRYAD, 2021) |
| *Entandrophragma candollei* | 0.644 | 9.885 | BR0000013947231 | RMCA (DRYAD, 2021) |
| *Entandrophragma cylindricum* | 0.647 | 11.35 | BR0000013990169 | RMCA (DRYAD) |
| *Erythrophleum suaveolens* | 0.876 | 13.7 | BR0000013649449 | RMCA (DRYAD, 2021) |
| *Euclinia longiflora* | 0.724 | 18.824 | 0 00000875246 8 | RMCA (DRYAD, 2021) |
| *Fernandoa adolfi-friderici* | NA | 12.744 | BR0000014165481 | NA |
| *Funtumia africana* | 0.489 | 12.551 | BR0000014262906 | RMCA (DRYAD, 2021) |
| *Garcinia kola* | 0.824 | 12.379 | BR0000016611474 | RMCA (DRYAD, 2021) |
| *Garcinia ovalifolia* | 0.825 | 10.227 | BR0000015735027 | RMCA (DRYAD, 2021) |
| *Garcinia punctata* | 0.802 | 8.371 | BR000001362192 | Harja et al. (2021) |
| *Garcinia quadrifaria* | NA | 12.807 | BR0000016607576 | NA |
| *Glyphaea brevis* | 0.669 | 14.847 | BR0000019969503 | RMCA (DRYAD, 2021) |
| *Greenwayodendron suaveolens* | 0.77 | 9.333 | BR000013690793 | Carsan et al. (2012) |
| *Grewia oligoneura* | 0.415 | 10.785 | 0 000006 87786 6 | Harja et al. (2021) |
| *Heisteria parvifolia* | 0.807 | 17.493 | BR0000019331010 | RMCA (DRYAD, 2021) |
| *Homalium africanum* | 0.701 | 8.96 | BR0000015975362 | RMCA (DRYAD, 2021) |
| *Hua gaboni* | 0.79 | 16.292 | 0 000008 71201 1 | RMCA (DRYAD, 2021) |
| *Hymenocardia ulmoides* | 0.777 | 5.98 | BR0000021708930 | RMCA (DRYAD, 2021) |
| *Irvingia excelsa* | 0.76 | 11.236 | BR0000013824440 | Harja et al. (2021) |
| *Irvingia grandifolia* | 0.948 | 9.962 | BR0000013610326 | RMCA (DRYAD, 2021) |
| *Klainedoxa gabonensis* | 0.946 | 8.113 | BR0000013613280 | RMCA (DRYAD, 2021) |
| *Lannea welwitschii* | 0.478 | 19.858 | BR0000013722050 | RMCA (DRYAD, 2021) |
| *Leplaea cedrata* | 0.536 | 10.901 | BR0000019201047 | Harja et al. (2021) |
| *Lepleae laurentii* | 0.563 | 17.389 | BR0000019201788 | Harja et al. (2021) |
| *Leptonychia tokana* | 0.591 | 7.647 | BR0000013680022 | RMCA (DRYAD, 2021) |
| *Lovoa trichiloides* | 0.635 | 9.452 | BR0000019204079 | RMCA (DRYAD, 2021) |
| *Macaranga monandra* | 0.477 | 19.802 | BR0000016376809 | RMCA (DRYAD, 2021) |
| *Macaranga spinosa* | 0.41 | 11.62 | BR0000020752286 | RMCA (DRYAD, 2021) |
| *Maesopsis eminii* | 0.48 | 25.115 | BR0000017141697 | RMCA (DRYAD, 2021) |
| *Magnstipula butayei* | NA | 7.652 | BR0000013351922 | NA |
| *Mammea africana* | 0.765 | 11.381 | BR0000013622770 | RMCA (DRYAD, 2021) |
| *Maranthes glabra* | 0.856 | 16.178 | BR0000015660831 | RMCA (DRYAD, 2021) |
| *Margaritaria discoidea* | 0.77 | 14.049 | BR000001441169 | Harja et al. (2021) |
| *Massulaira acuminata* | 0.869 | 9.394 | 0 00000 9 51205 4 | RMCA (DRYAD, 2021) |
| *Microdesmis yafungana* | 0.659 | 33.731 | BR00000133370008 | RMCA (DRYAD, 2021) |
| *Milicia excelsa* | 0.602 | 15.118 | BR0000018005509 | RMCA (DRYAD, 2021) |
| *Monodora angolensis* | NA | 36.172 | BR0000014057731 | NA |
| *Monodora myristica* | 0.528 | 5.079 | BR0000015300850 | RMCA (DRYAD, 2021) |
| *Musanga cecropioides* | 0.317 | 11.379 | BR0000018007480 | RMCA (DRYAD, 2021) |
| *Myrianthus arboreus* | 0.568 | 13.236 | BR0000013477592 | RMCA (DRYAD, 2021) |
| *Napoleonaea septentrionalis* | NA | 12.172 | BR0000015631510 | NA |
| *Nauclea diderrichii* | 0.695 | 9.23 | BR0000019765631 | RMCA (DRYAD, 2021) |
| *Nesogordonia kabingaensis* | 0.718 | 21.018 | BR0000013973933 | Harja et al. (2021) |
| *Oncoba welwitschii* | 0.704 | 9.041 | BR000001328297 | RMCA (DRYAD, 2021) |
| *Ongokea gore* | 0.787 | 12.165 | BR0000015994028 | RMCA (DRYAD, 2021) |
| *Pancovia harmsiana* | 0.802 | 11.698 | BR0000013987466 | RMCA (DRYAD, 2021) |
| *Pancovia laurentii* | 0.759 | 12.433 | BR0000015247650V | RMCA (DRYAD, 2021) |
| *Panda oleosa* | 0.607 | 12.226 | BR0000013565367 | RMCA (DRYAD, 2021) |
| *Paramacrolobium coeruleum* | 0.778 | 6.243 | BR0000016627871 | RMCA (DRYAD, 2021) |
| *Parinari excelsa* | 0.699 | 12.796 | BR00000177367233 | RMCA (DRYAD, 2021) |
| *Paropsia guineensis* | 0.855 | 3.539 | 0 000009 12887 3 | RMCA (DRYAD, 2021) |
| *Pentaclethra macrophylla* | 0.861 | 21.836 | BR0000018853162 | RMCA (DRYAD, 2021) |
| *Pericopsis elata* | 0.753 | 8.075 | BR0000013588564 | RMCA (DRYAD, 2021) |
| *Petersianthus macrocarpus* | 0.681 | 19.911 | BR0000013200152 | Harja et al. (2021) |
| *Phyllocosmus africanus* | 0.782 | 15.396 | BR0000016431492 | RMCA (DRYAD, 2021) |
| *Piptadeniastrum africanum* | 0.725 | 9.416 | BR0000018855456 | RMCA (DRYAD, 2021) |
| *Platysepalum chevalieri* | 0.85 | 6.102 | BR0000018332056 | Harja et al. (2021) |
| *Pleiocarpa pycnantha* | 0.681 | 10.593 | BR0000014550300 | RMCA (DRYAD, 2021) |
| *Prioria balsamifera* | 0.546 | 10.585 | BR0000013573249 | RMCA (DRYAD, 2021) |
| *Prioria oxyphylla* | 0.648 | 8.444 | BR0000013573812 | RMCA (DRYAD, 2021) |
| *Psydrax subcordata* | 0.609 | 7.901 | 0 00001256721 8 | RMCA (DRYAD, 2021) |
| *Pteleopsis hylodendron* | 0.72 | 9.669 | BR000001509065 | RMCA (DRYAD, 2021) |
| *Pterocarpus soyauxii* | 0.663 | 10.729 | BR0000013263430 | RMCA (DRYAD, 2021) |
| *Pterygota bequaertii* | 0.56 | 15.371 | BR0000013256852 | RMCA (DRYAD, 2021) |
| *Pycnanthus angolensis* | 0.551 | 20.316 | BR0000024584968 | RMCA (DRYAD, 2021) |
| *Quassia silverstris* | NA | 12.237 | BR0000019682662 | NA |
| *Ricinodendron heudelotii* | 0.354 | 15.746 | BR0000013987336 | RMCA (DRYAD, 2021) |
| *Rothmannia hispida* | 0.825 | 19.179 | BR0000021251078 | RMCA (DRYAD, 2021) |
| *Rothmannia macrocarpa* | NA | 17.387 | BR0000018694598 | NA |
| *Scaphopetalum thonneri* | 0.68 | 6.338 | BR0000018066937 | RMCA (DRYAD, 2021) |
| *Schumanniophyton magnificum* | 0.57 | 10.823 | BR0000013554675 | RMCA (DRYAD, 2021) |
| *Scorodophloeus zenkeri* | 0.655 | 6.977 | 0 00006 27433 7 | RMCA (DRYAD, 2021) |
| *Scytopetalum pierreanum* | NA | 11.307 | BR0000018120851 | NA |
| *Staudtia kamerunensis* | 0.786 | 7.715 | BR0000013978556 | Harja et al. (2021) |
| *Streblus usambarensis* | NA | 19.626 | BR0000017940542 | NA |
| *Strombosia grandifolia* | 0.761 | 6.876 | BR0000013964429 | RMCA (DRYAD, 2021) |
| *Strombosia pustulata* | 0.767 | 9.797 | BR0000015997296 | RMCA (DRYAD, 2021) |
| *Strombosiopsis tetrandra* | 0.677 | 18.816 | BR0000015225771V | RMCA (DRYAD, 2021) |
| *Symphonia globulifera* | 0.713 | 9.652 | BR0000015548511 | RMCA (DRYAD, 2021) |
| *Synsepalum brevipes* | 1.034 | 14.39 | BR0000019415031 | RMCA (DRYAD, 2021) |
| *Synsepalum stipulatum* | 0.741 | 5.804 | BR0000019421896 | RMCA (DRYAD, 2021) |
| *Synsepalum subcordatum* | 0.76 | 5.616 | BR0000019411927 | RMCA (DRYAD, 2021) |
| *Tabernaemontana crassa* | 0.585 | 8.69 | BR0000014341700 | RMCA (DRYAD, 2021) |
| *Tessmannia africana* | 0.75 | 16.142 | BR0000021547508 | RMCA (DRYAD, 2021) |
| *Tessmannia anomala* | 0.836 | 6.733 | 0 000009 88703 9 | RMCA (DRYAD, 2021) |
| *Tetrapleura tetraptera* | 0.726 | 9.303 | BR0000016148314 | RMCA (DRYAD, 2021) |
| *Tetrorchidium didymostemon* | 0.449 | 16.785 | BR0000016237407 | RMCA (DRYAD, 2021) |
| *Treculia africana* | 0.625 | 14.329 | BR0000017940900 | RMCA (DRYAD, 2021) |
| *Tricalysia coriacea* | NA | 6.968 | BR0000019622866 | NA |
| *Trichilia gillettii* | 0.597 | 11.732 | BR0000019211183 | RMCA (DRYAD, 2021) |
| *Trichilia prieureana* | 0.668 | 14.489 | BR0000013015251 | RMCA (DRYAD, 2021) |
| *Trichilia rubescens* | 0.557 | 12.468 | BR0000019216119 | RMCA (DRYAD, 2021) |
| *Trichilia welwitschii* | 0.58 | 15.084 | BR0000019218625 | RMCA (DRYAD, 2021) |
| *Tridesmostemon omphalocarpoides* | 0.782 | 8.247 | BR000001914676 | RMCA (DRYAD, 2021) |
| *Trilepisium madagascariense* | 0.499 | 28.334 | BR0000013570606 | Harja et al. (2021) |
| *Turreanthus africanus* | 0.5 | 13.601 | BR0000013975265 | Harja et al. (2021) |
| *Uapaca guineensis* | 0.712 | 20.062 | BR0000017450331 | RMCA (DRYAD, 2021) |
| *Vangueriella orthacantha* | 0.727 | 27.368 | 0 00000 9 02617 9 | RMCA (DRYAD, 2021) |
| *Vernonia conferta* | NA | 7.239 | BR000001494128 | NA |
| *Vitex ferruginea* | NA | 8.368 | BR0000018426083 | NA |
| *Voacanga africana* | NA | 19.003 | BR0000014680915 | NA |
| *Xylopia aethiopica* | 0.646 | 14.254 | BR0000014585104 | RMCA (DRYAD, 2021) |
| *Xylopia phloiodora* | NA | 11.23 | BR0000014129766 | NA |
| *Xylopia staudtii* | 0.498 | 11.882 | BR0000014130816 | RMCA (DRYAD, 2021) |
| *Zanthoxylum gilletii* | 0.646 | 27.513 | BR0000019428314 | RMCA (DRYAD, 2021) |
